# High gene dosage changes transcriptional regulation of genetic structures and contributes to diversity and heterogeneity

**DOI:** 10.1101/505446

**Authors:** Dianbo Liu, Ryan Hannam, Luca Albergante

## Abstract

Differences in gene dosages between and within species are widely observed. Nevertheless, our understanding of the impacts of gene dosages on transcriptional regulation is far from complete and it is largely ignored in many genetic studies. In this article, we showed that dynamic properties of three genetic structures became significantly different when gene dosages were high and, for self-inhibiting genes, high gene dosages could facilitate monoallelic expression at single cell level and heterogeneity of cell population. This shed lights on potential links between levels of ploidy and phenotypes. In addition, in the context of modern molecular biology, especially synthetic biology, gene dosage needs to be taken into consideration when designing or modifying genetic circuits.

## Introduction

Transcriptional regulation is the key control mechanism of gene expression which is required to regulate biological activities such as replication, stress response or apoptosis. Extensive scientific efforts were put to understand this process both at the molecular level, e.g., by looking at the effect of genetic mutations or changes on the structures of molecular complexes, and at the systems level, e.g., by studying the properties of multi-gene regulatory circuits such as transcriptional cascades or feedback loops [1, 2]. However, most of these analyses regarded genes as unique entities, ignoring how the copy number of particular gene(s) in a single cell, or called gene dosage, affected the dynamics of gene expression. This is potentially limiting as an increase in gene dosage can affect global gene expression either by increasing the RNA concentration, due to more “parallel” transcription, or by decreasing the nuclear concentration of transcription factors binding to that genes, due to an increased competitive binding.

Considerable levels of intraspecific gene dosage variation among individual cells were recently reported in multiple species including human and the effects of gene dosage variation were also supported by biological evidence [3, 19]. In bacterial cells, increased gene dosage was observed during the stationary phase of growth and this phenomenon was suggested to be related to regulation of gene expression [1]. One example in human is that of chromosomal trisomy, in which gene dosage of the whole chromosome increases by half, and is generally associated with severe phenotypes (e.g., Down syndrome) and is usually lethal. Even when a gene is duplicated, genetic effects can be observed, for example, an NFAT dysregulation in human was found to be induced by an increased dosage of DSCR1 and DYRK1A [2]. Hyperploidy was also often observed in human cancer [3, 4] and it was suggested that the presence of multiple gene copies might be instrumental for cancer cells to activate alternative genetic programs [5]. Although in mammals gene dosage variation is generally associated with pathological conditions, in other organisms this is not always the case. For example, dosage variation was believed to be important for the survival of different organisms, including prokaryotes [6], eukaryotic cells infected by viral genomes [7], yeasts [4], the parasite *Trypanosoma brucei* [8], plants[9], and enteric parasites such as *Entamoeba* [7, 10].

To investigate the effects of gene dosage variation at systems level, we decided to approach this issue by employing mathematical analysis of different genetic circuits that are found to recur in genetic networks across different organisms, i.e., self-inhibition, cross-repressors and inhibition chains [11-13]. In particular, we explored the changes in the quantitative dynamics of gene expression of the circuits when the gene dosage increased from low (≈ 1) to very high (≫ 1). Our findings, which are supported by deterministic and stochastic modelling of gene dynamics, indicated that the circuit dynamics was significantly affected by the gene dosages and hence that the gene dosage was an additional dimension that needs to be considered when inferring the behaviours of a genetic system from its topology. This finding shed light on the evolutionary reasons that potentially lead certain organisms to strongly change their gene copy number during specific situations. Moreover, it will be helpful to take gene dosage into account when interpreting the data obtained from synthetic biology experiments, as the designed biological circuits are often present in various numbers of copies.

## Results

### Interspecific and intraspecific differences in gene dosages are observed across multiple domains of life

Interspecific and intraspecific changes of gene dosages, are observed in different domains of life, including eukaryotes, prokaryotes and viruses. In evolution, changes of number of ploidy, thus changes in gene dosages, are widely observed. It was estimated that about 15% of angiosperm and 31% of fern speciation are accompanied by ploidy increase [14] and within the genera *Botrychium,* ploidy levels vary from 2N to 6N [15].

Intraspecific gene dosage changes occur through cell cycles, changes of growth conditions and in other situations. One example in eukaryotes is in members of foraminifera which are a phylum of amoeboid protists. Ploidy level of foraminifera can change from 1N to 100N during its nuclear cycles [7]. In human, the ploidy level is 1N in sperm and oocyte and was reported to be up to 16N in certain mature somatic cells in liver [16]. As for prokatyotes, DNA contents in *Escherichia coli* varies from 2 to 8 genome equivalents in stationary phase to up to 11 genome equivalents in exponential phase [1]. In addition to cellular organisms, the number of copies of genomes could vary significantly in the infected cells. For example, number of DNA viral genomes of podovirus P60 in a host cell can be up to 100 [17]. All these observed variations in levels of ploidy and gene dosages may affect behaviours of genetic circuits in the corresponding cells.

### High dosages of self-inhibiting gene facilitate monoallelic expression and phenotypic diversity

Negative feedback is a widely observed regulatory mechanism in natural and artificial systems, and is found in different biological processes such as transcription, translation and protein signalling. This mechanism imbues the system with properties such as an ultrasensitive response or short rising time [18]. We explored whether, with a negative feedback, a gene with high dosage in the genome will show different expression profiles compared with single copy. In this project, we only focus on transcriptional regulation but the results can be generalised to other regulatory mechanisms with similar properties. A demonstration of self-inhibiting gene is shown in Figure 1A. There are one or more identical genes and their constitutive promoters. All these gene copies produce the same products which bind to all the promoters and inhibit expression of the corresponding gene copies.

**Table 1.** Examples of variations of ploidy levels across and within species

| Level of ploidy / equivalent ploidy |  |  |  |  |
| --- | --- | --- | --- | --- |
| across species | within species |  |  |  |
| *Botrychium* genera | Human | foraminifera | *Escherichia coli* | podovirus |
| 2-6N | 1-16N | 1-100N | 2-8N | 1-100N |

**Figure 1.**
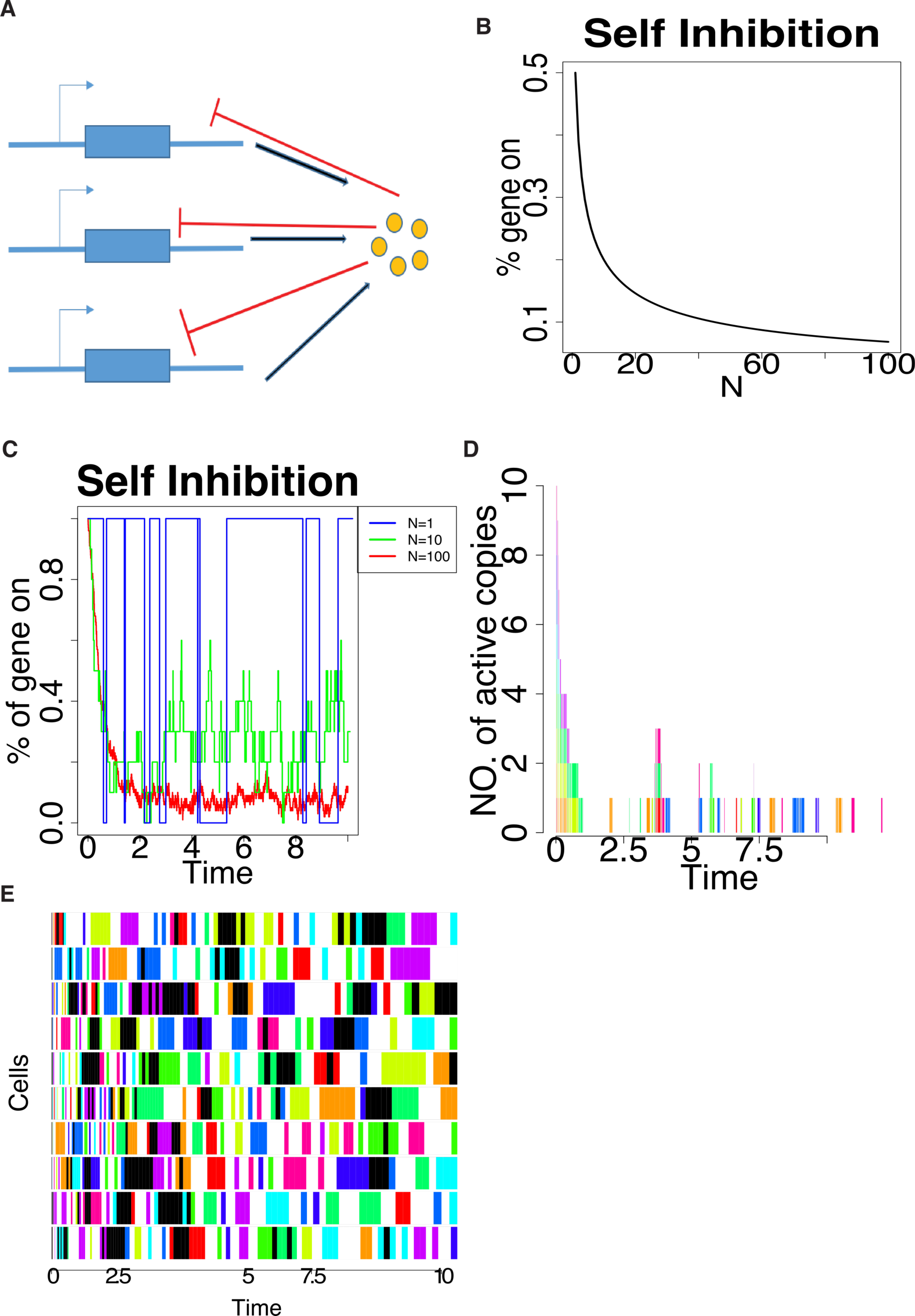
Self-inhibiting gene. (A) Self-inhibiting gene with gene dosage N >1. One or more copies of the same gene and their promoters are in a single cell. All these copies produce the same products which bind to the promoters and inhibit expression of all copies. (B) Percentage of actively expressing (uninhibited) alleles at steady state decreased as gene dosage increased. This phenomenon was captured using deterministic model (C) Stochastic model showed changes of percentage of actively expressing alleles over time at different gene dosages. Consistent with deterministic model, genes with a high dosage had a low percentage of actively expressing copies after initial period. (D) When the self-inhibition was strong, after initial stage, only a single (or a low number of) copies was on and the active copy changed from time to time. Different colours indicated different copies being expressed. Gaps means all the alleles are inhibited. (E) In cell population in the condition of (D), the self-inhibition system brought heterogeneity and thus, diversity, in the cell population. Each row is a single cell in the population, different colour indicates different copies of the same gene of interest being expressed. White colour means no allele was active and black indicates there are more than one alleles being simultaneously expressed in a single cell. The actively expressing allele switched over time in each individual cell and a wide range of different copies were expressed in the population. If these copies were slightly different from each other but sharing identical regulatory mechanisms, this mechanism gives diversity in the population.

A mathematical modelling approach was used to describe the system. A simple deterministic representation of gene A with *N* copies in each cell and with self-inhibition can be demonstrated as equations below:

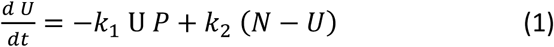

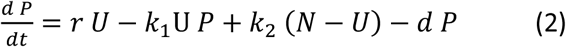

In the equation 3.1 and 3.2 above, *N* is the number of copies per cell (gene dosage) for the self-inhibiting gene. U stands for number of active copies (unbound by the products) and P is the number of products in cell. There are four reaction constants, *r* for net expression rate (transcription and translation), *d* for degradation rate of products, *k*_1_ and *k*_2_ for binding/unbinding rates respectively of the products to the promoter(s). This system has a single steady state. *U*′ and *P*′ are the number of active copies and number of products at steady state (Equation 3.3 and 3.4). While the level of products increased with gene dosage, the percentage of gene being actively expressed decreased with dosage (Equations 3.3-3.5 and Figure 1B).

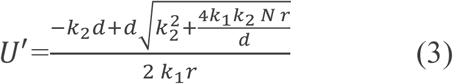

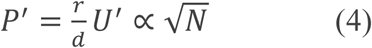

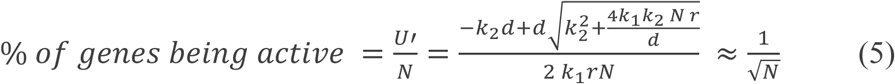

In addition to the deterministic model, the Gillespie algorithm was used to explore stochastic behaviours of the system and it showed similar behaviour, i.e. on average, lower proportion of genes are on when total number of gene copies is high (Figure 1C). More importantly, the stochastic model was able to capture some behaviours that could not be detected using the deterministic method. One interesting finding was, in each single cell, when the number of copies of the self-inhibiting gene was high (eg. 20), the system was able to establish and maintain single copy (or low number of copies) expression with frequent switching among alleles (as shown in Figure 1 D, each colour represents a different gene copy). If the expressed products of alleles were slightly different from each other in their structures and biological properties while still sharing the same regulatory machineries, this multiple-copy-self-inhibition system enables cells to switch among alleles and, therefore, phenotypes. At the population level, it can also be showed that frequent switches among alleles is able to facilitate diversity of phenotypes in the population while keeping the singularity (i.e. expression of a single allele) of each cell (Figure 1E).

### Cross-inhibiting genes with high dosages make a sharp tunable genetic switch

Cross-inhibiting gene pairs are frequently used and well-studied genetic circuits [19, 20]. We demonstrated that behaviours of cross-inhibiting genes changed considerably as dosages of both of the genes varied. In this system, gene A and gene B express repressors which bind to the promoter of the other gene and inhibit its expression but not their own promoters (Figure 2 A). The effect of gene dosages on the transcriptional profiles of the system was explored using both deterministic and stochastic models assuming both genes have the same dosage (*N*_*a*_ = *N*_*b*_ = *N*). Behaviours of the transcriptional cross-inhibiting genes can be modelled in a deterministic way using the following equations.

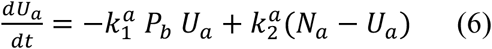

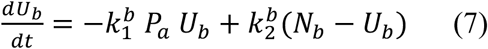

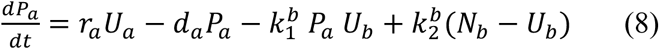

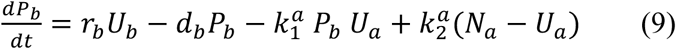

**Figure 2.**
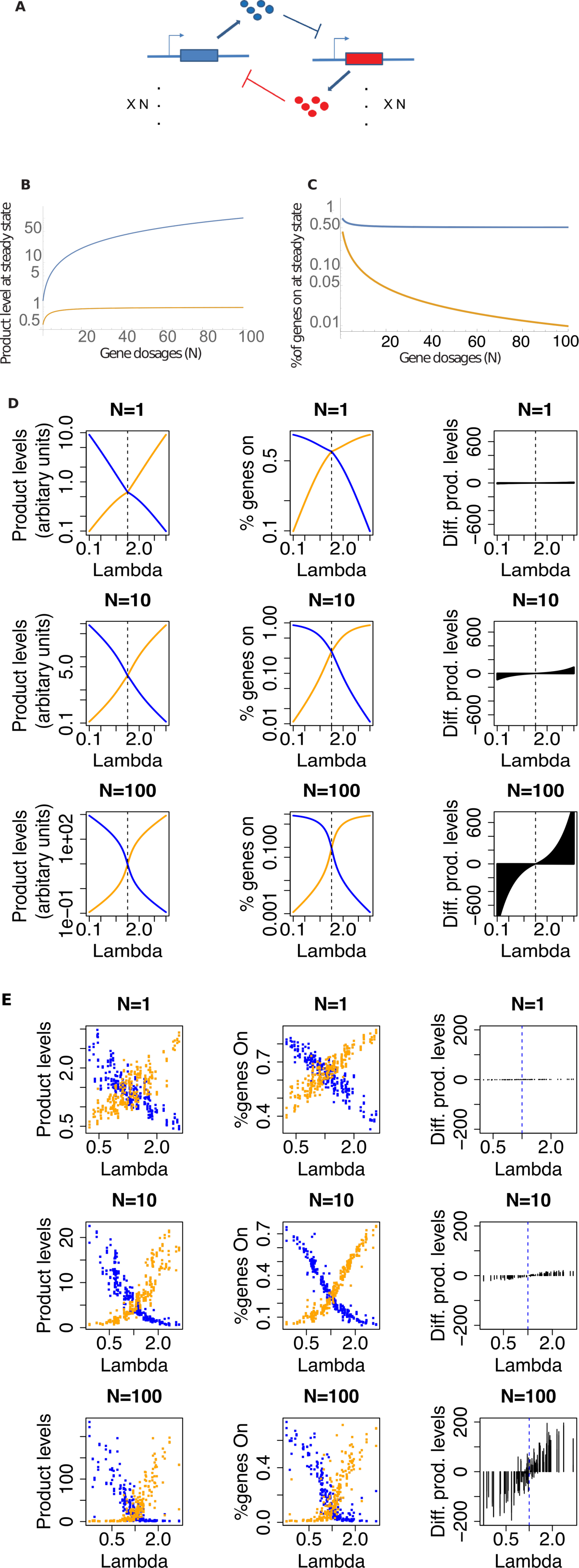
Cross transcriptional repressors. Analyses in B-D were conducted using the deterministic model and simulation in E was conducted using a stochastic model. (A) In this system, gene A and gene B expressed repressors to inhibit each other by binding to the corresponding promoters. The gene dosages (N) of A and B were the same and ≥ 1. (B, C) when one of the two repressors was dominating, the bigger the gene dosages were, the larger the difference there was between product level or percentage of actively expressing alleles at steady state. (D) Relative levels of the two repressors and percentage of active copies of each gene varied as the super parameter lambda changes. The levels of the two repressors converge when *λ* equals 1. The bigger the gene dosage, the larger the differences were between product levels of the two genes at steady state. (E) Same behaviours were observed using stochastic model.

In equations 3.6-3.9, *N*_*a*_ and *N*_*B*_ are gene dosages of gene A and B respectively. *U*_*a*_ and *U*_*B*_ are the numbers of active copies (unbound by the products) per cell. *P*_*a*_ and *P*_*B*_ are the numbers of gene products per cell. There are eight reaction constants, *r*^*a*^ and *r*^*b*^ for net expression rate (transcription and translation), *d*_*a*_ and *d*_*B*_ for degradation rate of products, 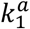 and 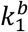 for binding rates of the products to the promoter(s) and, 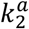 and 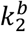 for unbinding rates of the products from the promoter(s).

In this project, we did not take into consideration cooperative binding, which will be discussed in follow-up research, and thus the Hill coefficient=1. This system had a single nontrivial steady state (see Materials and Methods). When all other parameters are fixed, the difference between the product levels of the two genes at steady state grows larger as the gene dosage (*N*) increased. (Figure 2B). The percentage of dominating gene being actively expressing (at steady state) was constantly high but that for the recessive gene kept decreasing as *N* became larger (Figure 2C).

Next, we analysed the equation to search for parameters that were likely to affect the relative expression levels of gene A and B. Amazingly, the relative expression level of the two genes only depended on a single summing-up ‘super’ parameter λ which is the ratio between all the eight parameters (Equation 3.10). When λ shifted from < 1 to > 1, there was a sharp switch between relative expression levels and percentage of active copies of the two genes. When λ = 1, expression level and portions of active copies of the two genes converge (Figure 2D). The difference between expression levels of dominating and recessive genes at steady state increased as λ deviated more from 1. The switch as λ passed 1 was sharper when *N* was large (>>1). A Gillespie algorithm was utilised to analyse the stochastic behaviours of the system and similar behaviours were observed (Figure 2 E).

Let:

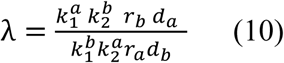

At steady state, when *N* = *N*_*b*_ = *N*_*a*_was large:

If λ=1

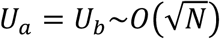

If λ>1

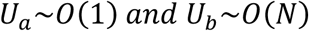

If λ<1

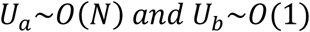

This finding suggested the balance of expression between the two cross-inhibiting genes could be adjusted by shifting the super parameter λ. For example, λ can be changed by altering the expression rate of the genes (parameter *r*_*a*_ or *r*_*b*_), for example by introducing an additional inducer to inhibit expression of one gene in the system. This multi-copy-cross-inhibiting system can be used as a tunable genetic switch and deliver sharp responses when *N* is high and provides a simple and flexible options to construct sharp switches [21].

### High gene dosages decrease the attenuating effects of long linear inhibition chains

Long linear regulatory chains (LRCs) is an interesting structure in genetic networks[11, 12, 22, 23], in our previous publication, we showed, in gene regulatory networks, LRCs reduced the effect of the regulation from input signals by bringing down relative effectiveness (RE) and thus make long LRCs inefficient in passing information (Figure 3 A and B) [11, 12]. An example of an inhibitory LRC, in which each gene is inhibited and only inhibited by its direct upstream gene via transcription factor, is shown in Figure 3A. Definition of RE is given in equation 3.11 and 3.12.

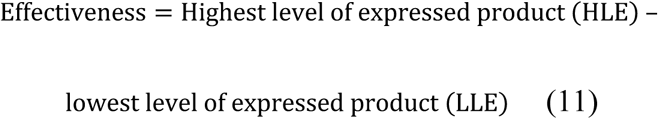

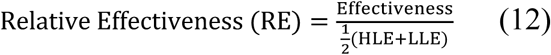

In this article, RE attenuation effects of LRCs was found to be weakened if dosage (*N*) of each gene in the chain was large (>>1). Transcriptional regulation was described using the following equations (only the first three genes in an LRC are shown) :

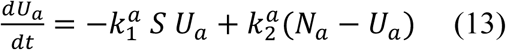

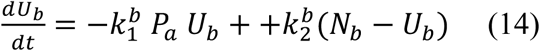

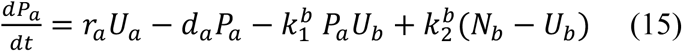

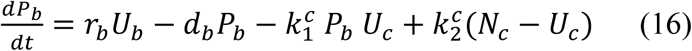

**Figure 3.**
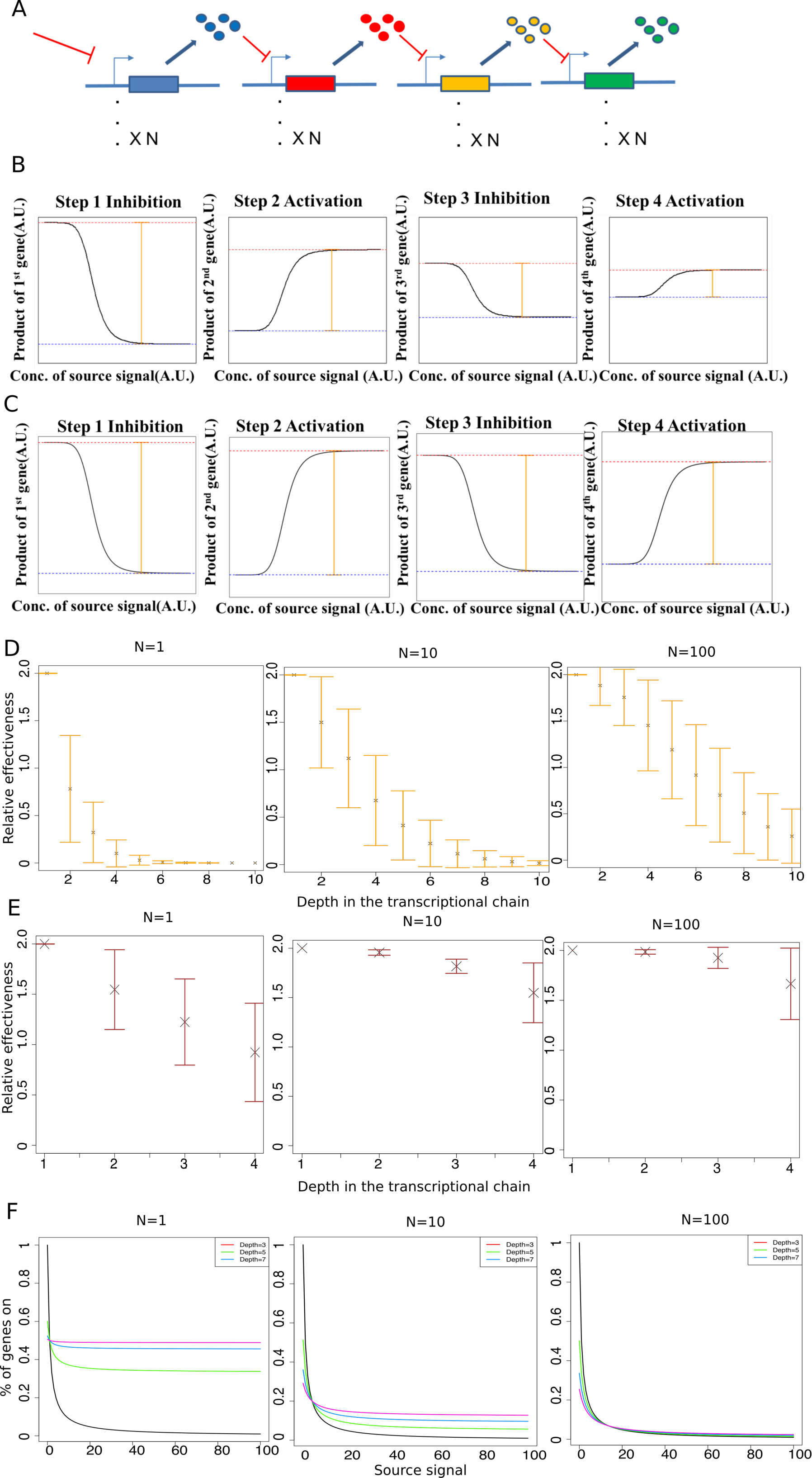
Transcriptional inhibition chain. (A) Inhibition chains are linear chains of one-way regulation in which each gene inhibits its downstream gene via its product as a repressor. There is an external source inhibitory signal acting only on the first node in the chain. (B) An example of how effectiveness (orange) decrease along the chain. (C) In an inhibition chain with large gene dosages (N) for each node, relative effectiveness decreases slower. (D) Trend of decrease of relative effectiveness was attenuated when dosage of each gene was high in the chain. (E) Stochastic simulation showed similar results as deterministic calculation. (F) Percentage of genes on at odd depths of the chain (net inhibitory) decreased as gene dosages increased.

In equations 3.13-3.16, *N*_*a*_, *N*_*B*_ and *N*_*C*_ are gene dosages of gene A, B and C respectively, where products of gene A binds to promoter of gene B and inhibits expression of gene B and products of gene B inhibit gene C. *S* is the external signal that inhibits expression of gene A, for example, a chemical that binds to the promoter of gene A and inhibits its expression. *U*_*a*_, *U*_*B*_ and *U*_*C*_ are the numbers of active copies (unbound) per cell. *P*_*a*_ and *P*_*b*_ are the numbers of gene products per cell. There are eight reaction constants, *r*_*a*_ and *r*_*b*_ for net expression rate (transcription and translation), *d*_*a*_ and *d*_*b*_ for degradation rate of products, 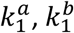 and 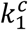 for binding rates of the products to the promoter(s) and, 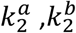 and 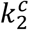 for unbinding rates of the products from the promoter(s).

The magnitude of RE attenuation went down as dosage of all genes increased assuming dosages were the same for all the genes (Figure 3D). In addition, at odd (net inhibitory) depth of the LRCs, percentage of genes that were active (unbound by inhibitors) decreased as gene dosage increased. This observation suggested longer LRCs can be more efficient in signal transduction in transcriptional networks of organisms with higher ploidies and, when designing genetic circuits in synthetic biology, long LRCs could be functional if the gene dosage is high. A stochastic model was also applied to analyse the system and showed similar results (Figure 3 E). This observation suggests that LRCs could be more abundant in cells with high gene dosages or higher number of ploidy because they are more efficient in passing signals, which is consistent with our previous observation that in human, gene regulatory chains are shorter in human non-cancer cell line compared with cancer cell line[11, 12].

## Discussion

Genomes are traditionally perceived as having fixed gene dosages for most genes within many species, especially eukaryotes, and thus individuals within the same species are thought to share nearly identical genomes. However, this static notion of the genome was challenged by a variety of studies in recent years and the dynamic nature of genomes was shed light on [7]. In eukaryotes the deviation from ‘standard’ versions of haploid-diploid by nuclear cycles can facilitate significant changes in karyotypes and thus global gene dosages. In prokaryotes and in viral infections, dramatic gene dosage changes are also observed under different growth phases or environmental conditions. In this article, transcriptional dynamics of three different genetic circuits were studied and shown to be different between low and high gene dosages. These findings suggested potential changes in biological activities associated with a shift in gene dosages. In evolution, these qualitative differences in transcriptional regulation might also be one of the reasons why most duplicated genes are eventually lost following whole genome duplications [24, 25]. Within a species, variation in genome contents and gene dosage could be a widespread source of phenotypic variation [7].

In addition to transcriptional regulation, gene dosage can affect functions of cells in multiple biological processes. Gene products interact with other cellular molecules and form complexes. If stoichiometric balance is broken by variation of gene dosage, the fitness of a cell may decrease [26], which has been pointed out in aneuploidy of eurytopic cells [27]. Imbalances of gene products caused by gene dosage variation were also reported to have impact on functions of signaling pathways such as mitogen-activated protein kinase signaling pathway, where changes of gene dosage was suggested to modify response patterns [4, 28, 29]. In this project, we demonstrated that gene dosages changed behaviors of transcriptional circuits and potentially phenotypic differences at both single and population levels. To have a complete understanding of cellular effects of gene dosages, all these aspects need to be taken into consideration.

Multiple mechanisms at transcriptional, post-transcriptional and protein-protein interaction level were suggested to buffer the effects of gene dosage changes in cells. Veitia *et al.* suggested that, at transcriptional level, the origin of the buffering effect came from the fact that transcription factors have different affinities to its binding sites in cis-regulatory regions [26]. At post-transcriptional level, production of rapidly degraded transcripts was thought to be related with gene dosage variations. At protein level, It was shown theoretically that multimeric proteins and preferential protein degradation were relevant in gene dosage effects [30, 31]. It is worth mentioning that many of the buffering or compensation mechanisms are related to the nonlinear nature of the biochemical reactions which is however a large topic and is beyond the scope of this project. Nevertheless, it could be possible, according to our results and findings in literature, that gene dosages, or number of ploidy, and nonlinearity of biochemical reactions form two different lines of evolutionary strategies, potentially at different stages of evolution, one to implement quick behavioral changes while the other improving efficiency of energy and material usage. Follow-up research work will be conducted to specifically analyze the effects of gene dosage on dynamic behaviors of the genetic circuits while taking nonlinearity into consideration.

## Materials and Methods

### Code and simulation

All the simulations, calculations and generations of graphs were conducted under R environment (version 3.2.3.). The codes are available at https://github.com/kaiyuanmifen/GeneDosage

### Self-inhibition

The levels of products and percentages of active gene copies (Figure 1) were calculated according to equations presented in Results section. The parameters used are *k*_1_ = 1, *k*_2_ = 1, *r* = 2, *d* = 1 and *N* = 1: 100. The stochastic simulation was conducted using Gillespie algorithm based population of each species using the same parameters. The simulation to explore frequent changes of active gene copies and potential heterogeneity generated (Figure 1 E and F) were conducted using the same Gillespie algorithm with 10 copies of the self-inhibiting gene in each cell with parameters: *k*_1_ = 1, *k*_2_ = 1, *r* = 2, *d* = 1.

### Cross-repressors

With no cooperative binding (Hill coefficient =1), the system we modelled has a single nontrivial steady state [19]. The product levels and percentages of active copies were calculated according to the steady state. Parameters for the analysis in Figure 2 were *k*_12_ = 1, *k*_21_ = 1, *r*_1_ = 2, *d*_1_ = 1, *v*_12_ = 1, *v*_21_ = 1, *r*_2_ = 1, *d*_2_ = 1 with dosages (*N*) for both genes varying from 1 to 100. For analysis in Figure 2 D and E, while *r*_1_ varied from 10 to 1 (*r*_2_ was fixed at 1) and *r*_1_varied from 1 to 10 (*r*_1_was fixed at 1) to give a range of 0.1 to 10 for lambda. The stochastic simulation was conducted using the same home-made Gillespie algorithm. Simulation was conducted with *N* = 1, 5, 10, 20, 30, 40, 50, 60, 70, 80, 90, 100 (only 1, 10 and 100 are shown). A total of 2000 rounds of simulation were conducted. Other parameters were randomly sampled using Latin-hypercube method within the range *k*_12_ in (1,2), *k*_21_ in (1,2), *r*_1_ in (2,4), *d*_1_ in (1,2), *v*_12_ in (1,2), *v*_21_ in (1,2), *r*_2_ in (2,4), *d*_2_ in (1,2) and λ was calculated accordingly. The differences between product levels of each gene was calculated using the mean levels from the simulations. Stability of the single steady states was analysed using Jacobian matrix. The steady state could be unstable (at least one out of four eigenvalues can be positive).

### Inhibition chain

The deterministic analysis of inhibition chain was conducted using equations presented in Results section. Parameters for each step of the chain was randomly sampled using Latin Hypercube method within the range: *k*_12_ in (0.1, 10), *k*_21_ in (0.1, 10), *r* in (0.1, 10), *d* in (0.1, 10). RE was calculated when source inhibitory signal varied from 0 to 10000. 1000 rounds of simulations for *N*= 1, 10 and 100 were conducted for chain of length 10 and standard deviation was calculated from the simulations. The same parameter sets were used for stochastic simulation, using Gillespie algorithm, based on population of each species with 30 replications were conducted for *N* =1,10 and 100. Four instead of ten nodes were simulated because of limitation of computational power. Percentage of active copies was calculated based on equations in results section. The parameters used were *k*_12_=0.1, *k*_21_=0.1, *r* =0.1, *d* =0.1 for all the depths.

